# Identifying and Interpreting Subgroups in Health Care Utilization Data with Count Mixture Regression Models

**DOI:** 10.1101/488924

**Authors:** Christoph F. Kurz, Laura A. Hatfield

## Abstract

Inpatient care is a large share of total health care spending, making analysis of inpatient utilization patterns an important part of understanding what drives health care spending growth. Common features of inpatient utilization measures such as length of stay and spending include zero inflation, over-dispersion, and skewness, all of which complicate statistical modeling. Moreover, latent subgroups of patients may have distinct patterns of utilization and relationships between that utilization and observed covariates. In this work, we apply and compare likelihood-based and parametric Bayesian mixtures of Negative Binomial and zero-inflated Negative Binomial regression models. In a simulation, we find that the Bayesian approach finds the true number of mixture components more accurately than using information criteria to select among likelihood-based finite mixture models. When we apply the models to data on hospital lengths of stay for patients with lung cancer, we find distinct subgroups of patients with different means and variances of hospital days, health and treatment covariates, and relationships between covariates and length of stay.

## 1 Introduction

Recent policy attention has focused on the triple aim of improving health outcomes and care quality while reducing health care spending. These efforts require a detailed understanding of the drivers of variation in spending and outcomes to target payment and delivery reform incentives. Inpatient hospital services account for the majority of total health care spending. [1] In this study, we wish to understand variation in hospital inpatient days among patients diagnosed with lung cancer. Lung cancer is the most common cancer worldwide and a major cause of cancer-related mortality. [2]

Health care utilization data such as days in the hospital or patient-level spending are nonnegative and often right-skewed, heavy-tailed, and multi-modal with a point mass at zero. Models for these data must be flexible enough to accommodate these features and still produce interpretable, policy-relevant results. [3] Generalized linear models (GLMs) with exponential family distributions such as Poisson, geometric, and Negative Binomial can accommodate non-negative and right-skewed count variables. [4–6] To account for excess zeros, these count models can be augmented with zero-inflation [4, 7, 8] or hurdle components. [9] Additional flexibility such as multi-modality and over-dispersion can come from mixture models. [10]

Mixture regression models are well known (see reviews in [11, 12]) and may be called switching models [13] or latent class models. [14, 15] Mixtures of count distributions, such as Poisson [16] and Negative Binomial, [17, 18] may also be augmented with a hurdle component for excess zeros. [19] Mixture models can also link the mixture component probabilities to covariates. [3]

Our second motivation for using mixture models is to discover latent subclasses of individuals with distinct utilization patterns, which mixture regression models also accomplish. [20] We want to identify patient subgroups with different patterns of inpatient lengths of stay. [21, 22] Policymakers, payers, and clinicians seeking to improve care and reduce spending in these groups could design interventions tailored to subpopulations. Previous work has shown substantial heterogeneity in the patterns of health care utilization among patients with lung cancer. [23] In this application, both the mixture components and their parameters are of interest, as are the clusters of observations drawn from each component.^1^

A key challenge in fitting mixture models is determining the number of mixture components. Too many components will over-fit the data and impair model interpretation, and too few will be insufficiently flexible. The number of components can be decided *ex ante*, by choosing a convenient and interpretable number such as two or three, or *ex post*, by calculating models with different numbers of components and comparing their fit statistics, such as Akaike Information Criterion (AIC) [24] or Bayesian Information Criterion (BIC) [25], or likelihood ratio tests. [26] In a Bayesian approach, the number of components can be treated as a parameter and informed by both the data and prior information. [27] Algorithms to fit Bayesian mixture models are diverse. The earliest approaches used reversible jump MCMC [28, 29], to accommodate the changing model size as the number of components changes across iterations, and data augmentation [30]. More recently, maximum *a posteriori* estimation [31], variational inference [32], and alternative MCMC algorithms [33, 34] have been proposed.

In this paper, we define and compare two implementations of mixture models for zeroinflated count regression: maximum-likelihood-based finite mixture models (FMMs) and parametric Bayesian mixture models. Previous authors have also used likelihood-based [35] and Bayesian [36, 37] models for zero-inflated health care utilization outcomes (see review in [38]). Others have fit both likelihood-based [17, 39] and Bayesian mixtures models for two-part count regressions, including for health claims data. Still other authors have fit finite mixtures of count regressions. [40] Our contribution to this literature is an explicit comparison between maximum likelihood and parametric Bayesian mixture models for (zero-inflated) count data. We compare these two approaches’ ability to detect the true number of mixture components and estimate component parameters, as well as the practicalities of both approaches.

The rest of the paper is organized as follows. First, we detail the count regression models and their mixture implementations in Section 2. Section 3 outlines our real data and simulations studies. We present model checks in Section 4 and results in Section 5. Finally, we conclude in Section 6 with implications of the results and suggestions for future work.

## 2 Model Definitions

### 2.1 Negative Binomial Regression

The Negative Binomial distribution accommodates over-dispersion in count variables with a longer, fatter tail than the Poisson distribution. [41] [42] identified numerous parameterizations of the Negative Binomial; here, we use the definition in [17]. For *n* = 1, …, *N* observations and *d* = 1, …, *D* covariates, the data comprise an (*N*, *D*)-dimensional covariate matrix **X** with rows **x***_n_* and an *N*-vector of outcomes **y** = (*y*_1_, …, *y*_*N*_)^′^. For simplicity, we omit the subscript *n* in what follows. The density function for the NegBin(*y|µ*, *Ψ*) distribution is

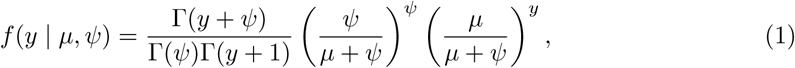

where *Ψ* is a precision parameter and we specify a regression model for the mean parameter

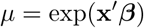

for covariate vector **x** = (*x*_1_, …, *x*_*D*_)^′^ and corresponding regression coefficient ***β*** = (*β*_1_, …, *β*_*D*_)^′^. In this specification, mean and variance are

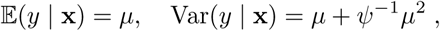

which corresponds to the NB2 model definition. [43]

### 2.2 Zero-Inflated Negative Binomial Regression

We extend the model above with *zero-inflation*. This models combines two sources of zeros: a point mass at zero and a Negative Binomial distribution, which generates both zero and non-zero count values. We can write a ZINB model *ZINB*(*y* | *µ*, *Ψ*, *π*) as

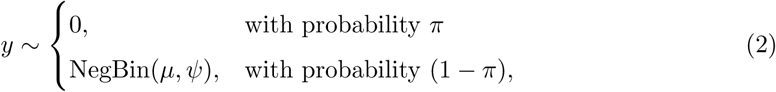

where *π* ∈ (0, 1) is the probability of an observation being a structural zero, and we model this probability with a Binomial distribution. Using a canonical log link, we specify a regression model for the probability

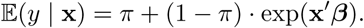

where **x** is a vector of covariates and ***β*** is a coefficient vector, as before.

### 2.3 Mixture Regression Models

The regression models detailed above can accommodate the zero-inflation, over-dispersion, and skewness of our count data. A mixture model implementation will allow us to discover latent subpopulations (i.e., clusters) in the data. Below we outline both likelihood-based and Bayesian mixture model implementations of (ZI)NB regressions.

### 2.4 Finite Mixture Models

Our first formulation is a finite mixture of *k* = 1, …, *K* Negative Binomial distributions, each parameterized by a mean *µ*_*k*_ = exp(**x*β****_k_*) and a precision parameter *Ψ*_*k*_. Let the contribution of each component to the mixture be denoted *c*_*k*_ such that *c*_*k*_ ∈ [0, 1] and 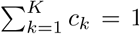. The distribution of the outcome *y* is the weighted sum over these Negative Binomial mixture components,

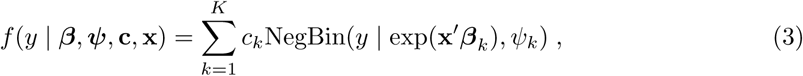

where 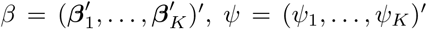, and **c** = (*c*_1_, …, *c*_*K*_)^′^. The extension to ZINB mixtures is straightforward with the addition of *π*_*k*_ parameters to govern the zero-inflation.

We add subscripts *n* = 1, …, *N* to identify individual observations of the outcome *y*_*n*_ and the covariates **x***_n_*. Then we augment these observed data with cluster membership indicators *z*_*nk*_, which equal 1 if observation *n* is drawn from component *k* and 0 otherwise. An observation can come from only one component, so ∑_*k*_ *z_nk_* = 1. The complete data likelihood is therefore

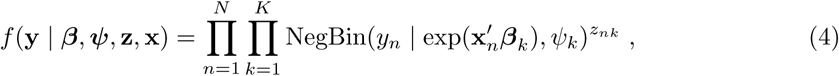

where **y** = (*y*_1_, …, *y*_*N*_)^′^ and 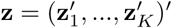 is the collection of *N*-vectors **z***_k_* = (*z*_1*k*_, …, *z*_*Nk*_)^′^.

Of course, the membership indicators for each observation are missing, because we do not know from which component any observation is drawn. Thus we use an expectation-maximization (EM) algorithm to fit the model. [44–46] The EM algorithm iterates between two steps to overcome the missing data problem. In each E step, we take expectation of the complete data log likelihood with respect to the conditional distribution of the missing component assignment indicators given the current parameter values and the observed data. By averaging over these for each component, we obtain prior probabilities of each component. Then in the M step, we maximize the expected log likelihood in the parameters.

In addition to mixing probabilities and parameter estimates for each component, the EM algorithm also produces posterior probabilities of each observation belonging to each component, i.e.,

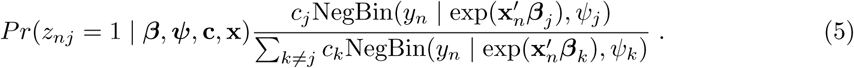

We can use the posterior probabilities to “hard classify” each observation into a cluster using the component for which it has the highest posterior probability of membership. Alternatively, we can weight observations by their posterior probabilities of being in each cluster. Using either method, the goal is to produce summaries of the clusters of observations.

The extension of the model above to mixtures of ZINB regressions is straightforward by replacing the Negative Binomial model in Eqs. (3) and (4) with the corresponding ZINB model,

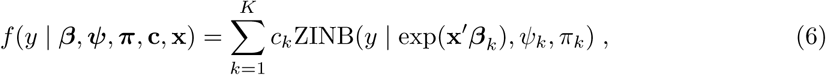

and

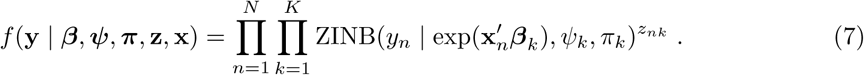

To fit the models in Eqs. (3) through (7), we modify the `FLXMRnegbin` function from the `countreg` package [47, 48], which is a driver for the general EM mixture model fitting algorithms of the `flexmix` package. [49–51] This package also implements stochastic EM (SEM), which uses a sample from the conditional distribution of the membership indicators rather than their expectation. To choose the number of mixture components *K*, we fit the model for various values of *K* and choose the one with the best fit statistics. We use AIC and BIC to compare models with different numbers of components.

### 2.5 Bayesian Mixture Models

We next turn to Bayesian mixture models, for which we specify a prior distribution on component membership indicators in Eq. (4). A popular choice is a multinomial distribution on the (binary, sum-to-one) indicators and a (conjugate) Dirichlet prior on the mixture probabilities.

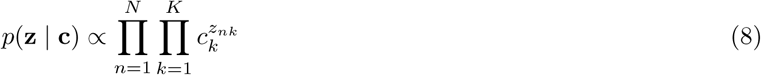

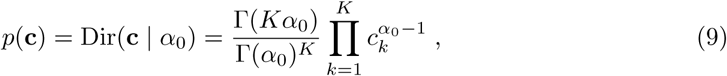

where Γ(·) is the Gamma function and *α*_0_ is a hyperparameter. We fix the maximum number of components at *K* = 20. In practice, the number of components with non-trivial posterior mixing proportions is less than *K*, resulting in mixture model that has only as many components as the data require. See Section 6 for more discussion of this choice.

To complete the specification, we need priors for the regression coefficients and precision parameters of the component regression models. We chose Normal and Log Normal distributions,

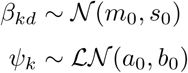

for *k* = 1, …, *K* and *d* = 1, …, *D* and hyperparameters *α*_0_ = 0.1, *m*_0_ = 0.0, *s*_0_ = 10.0, *a*_0_ = 0.0, and *b*_0_ = 2.0, that is, weakly informative priors. [52]

As before, this mixture model produces posterior estimates of the prevalence of each component and the parameters that govern it, plus posterior probabilities of membership in each mixture component for each observation.

The extension to mixtures of ZINB regressions follows as before, putting a multinomial-Dirichlet prior on the component indicators in Eq. (7). We refer to the models as MD-NB for the Multinomial-Dirichlet Negative Binomial and MD-ZINB for the zero-inflated version.

We implement this model in STAN [53] with 2000 iterations and a warm-up of 1000. Because this No-U-Turn sampler [54] does not support discrete latent variables, we marginalize over the component assignment variables.

## 3 Data

### 3.1 Monte Carlo Simulation

In a small simulation study, we compared the performance of the 2 modeling techniques described above: 1) finite mixture models with fixed numbers of components fit via EM and 2) Bayesian mixture models. For this, we generated data from Negative Binomial mixtures with 2, 3, 4, and 5 components and varying amounts of overlap. We first generated three covariates **x** = (*x*_1_, *x*_2_, *x*_3_)^′^ from Normal distributions *N* (0, 0.5). The corresponding intercept and regression coefficients ***β****_k_* = (*β*_0*k*_, *β*_1*k*_, *β*_2*k*_)^′^ and dispersion parameters *Ψ*_*k*_ were chosen to produce components with high, medium, and low overlap. Then we generated the outcome as *y* ∼ NegBin(exp(**x*β****_k_*), *Ψ*_*k*_). These choices (summarized in Appendix Table 9) produce a variety of shapes of the Negative Binomial mixture components with different degrees of overlap. Figure 1 shows densities of the true mixture components in each scenario.

**Figure 1:**
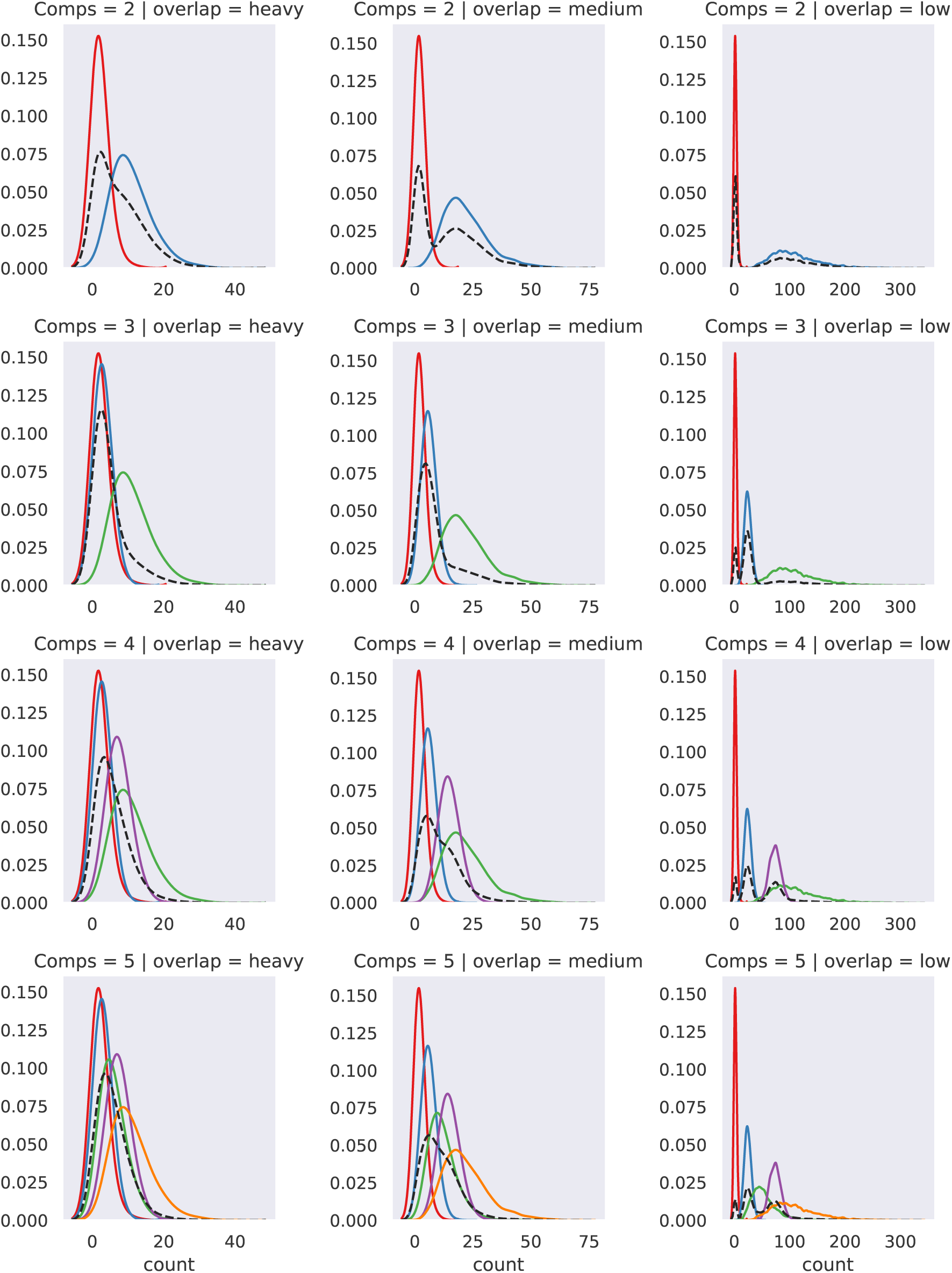
Density histograms for the simulated data sets with 2, 3, 4, and 5 mixture components with varying amounts of overlap. The black dashed line marks the combined density.

### 3.2 AOK Data Set

We analyzed health care billing claims provided by the AOK Research Institute. AOK covers around 30% of the German resident population. The data set contains patient-level information on inpatient and outpatient diagnoses and procedures from 2009 to 2012, as well as service utilization. We used a study population previously derived from this data set consisting of patients with incident lung cancer in 2009. More information on the data set and the selection criteria can be found in Schwarzkopf et al. [55]

The outcome of interest was the total number of inpatient hospital days for each patient in the year after diagnosis. Inpatient hospital days are defined as the number of days from formal admission to hospital until discharge (i.e., hospital outpatient procedures do not count as hospital days), summed over all hospitalizations in a year. Admission and discharge on the same day count as one hospital day, so only individuals who were never admitted have zero hospital days. We included only individuals who survived for the full year, resulting in 7118 individual observations. The mean number of hospital days is 44, with a maximum of 296 and 59 zero observations.

We included the following covariates in the model: age, sex, treatment type during the course of the disease (chemotherapy, radiation therapy, or surgery), number of other tumor sites at diagnosis, number of metastases at diagnosis, Charlson comorbidity index, and district type of residence (major city, urban district, rural district, or thinly populated rural district). The Charlson comorbidity index was calculated using ICD-10 codes as in [56] with the slight modification of excluding the diagnosis of lung cancer out of the group “solid tumor without metastases”.

Figure 2 summarizes the sample in a tableplot. [57] The skewness of the hospital days outcome is apparent in the leftmost panel, with wide variation at the highest quantiles of the distribution. Patients were mostly older than 60 years, male, and urban, and these demographic features did not appear to be strongly related to length of stay. Number of metastases, Charlson scores, chemotherapy, and surgery were all positively correlated with length of stay.

**Figure 2:**
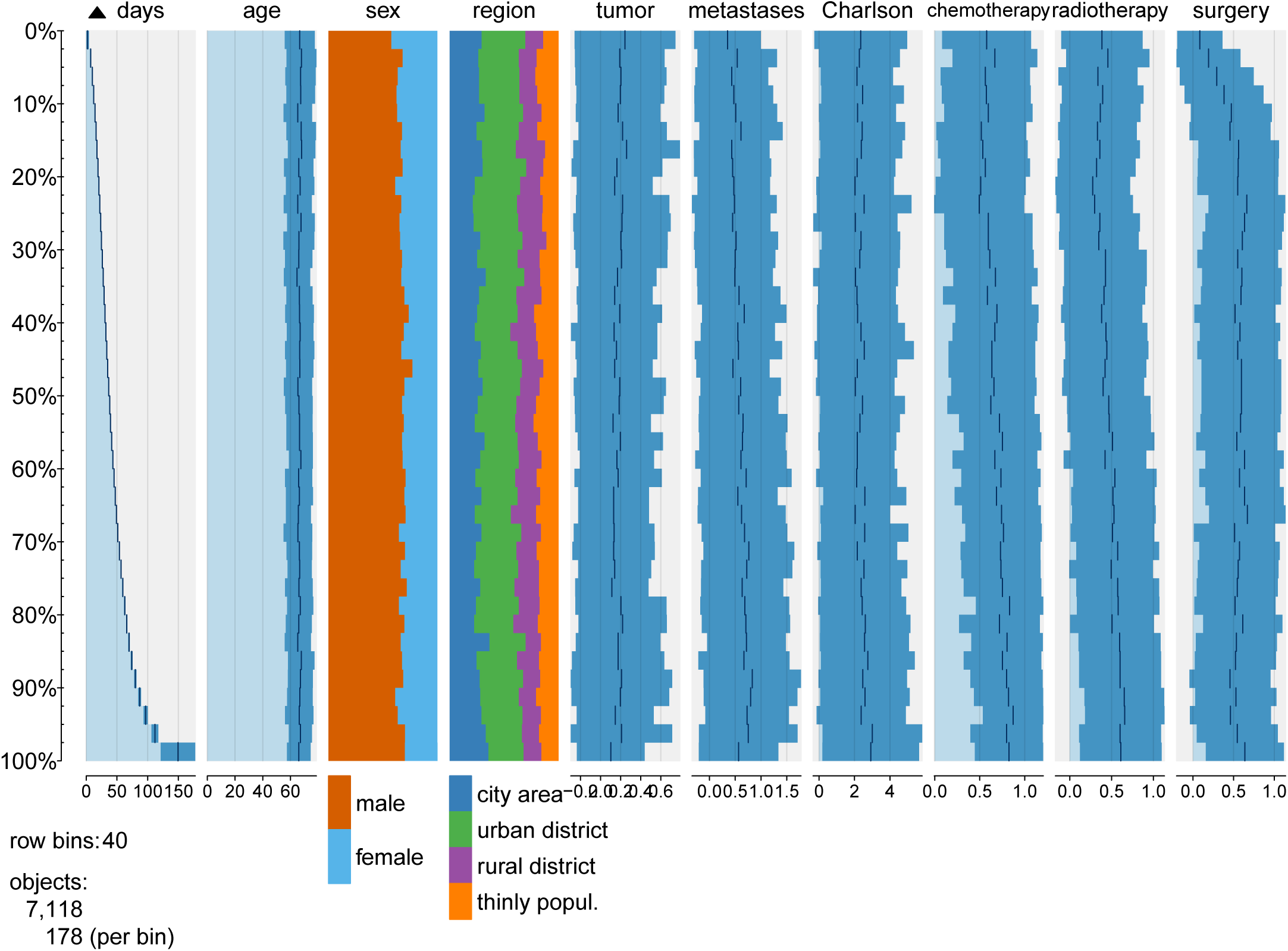
Tableplot of the AOK data set. The whole data set is sorted by hospital days (left side) in increasing order. All other variables are grouped into row bins, where numeric variables are displayed as bar charts and categorical variables as stacked bar charts.

## 4 Model Evaluation

### 4.1 Graphical Posterior Predictive Checks

To check the fit of the model, we use three different graphical checks. The first two are based on the posterior predictive, [58] which is the distribution of the outcome conditional on the posterior of the parameters, that is, after updating our beliefs about the parameters using the observed data. From the posterior predictive, conditional on the observed covariates **X**, we repeatedly simulate *N* observations (as in the observed data), 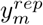 for *m* = 1, …, 1000. Then we compare the distribution of each 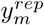 to the distribution of the observed outcome *y*. Similar shapes indicate good model fit.

We also compute a test quantity on each replicated data set, 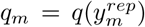, and plot the distribution of this test quantity compared with the observed test quantity *q* = *q*(*y*). Specifically, we use the mean as our test quantity of interest, 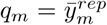. A distribution of simulated means centered around the observed mean supports good model fit.

Rootograms [59] show goodness-of-fit for count data by comparing observed frequencies from the model to expected frequencies. We can use these to check for over-dispersion, skewness, and excess zeros. We employ hanging rootograms in which the bars of observed frequencies are “hanging” from the curve representing the expected frequencies. Rootograms use “soft” component assignment weights, i.e., probabilities of belonging to a certain component. Ideal fit is indicated when the bars are close to the horizontal zero line.

## 5. Results

### 5.1 Component Identification in Simulated Data

Table 5.1 presents the results of the Monte Carlo simulation using the AIC, BIC, and MD-NB to estimate the number of components in the mixture.

AIC and BIC selection finds the true number of components in two of the simulations, for low and medium overlap with 4 components. On the other hand, MD-NB is very accurate in detecting the true number of components in all scenarios with 5 components. It also identifies the true 2 components for medium overlap and the true 4 components for high overlap.

All methods tend to overestimate the number of components in most scenarios but MD-NB is still closer to the truth. When the true number of components are estimated correctly, the regression parameter estimates and mixture weights are also close to the true values (see Table 9 and Table 9 in appendix).

### 5.2 Mixture Regression Analysis of AOK Data

For the AOK data set, the MD-NB finds 3 components as having the highest expected posterior mixture weights. In the following, we only show results based on this final model with 3 components. Figure 7 displays histogram plots of five replicated outcomes 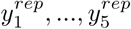 using the posterior predictive distribution. All replicates exhibit similar shape to the true distribution *y*. The means of the replicated data are centered around the mean of the observed data (Figure 8).

**Table 1:** True and estimated number of components based on AIC and BIC model selection criteria and on MD-NB for the simulated data sets. Bold indicates cases where the selected number of components matches the truth.

| Truth | high overlap |  |  | medium overlap |  |  | low overlap |  |  |
| --- | --- | --- | --- | --- | --- | --- | --- | --- | --- |
|  | AIC | BIC | MD-NB | AIC | BIC | MD-NB | AIC | BIC | MD-NB |
| 2 | 1 | 1 | 4 | 3 | 3 | **2** | 6 | 3 | 5 |
| 3 | 5 | 5 | 4 | 6 | 4 | 5 | 5 | 5 | 5 |
| 4 | 6 | 6 | **4** | **4** | **4** | 6 | **4** | **4** | 5 |
| 5 | 6 | 6 | **5** | 4 | 4 | **5** | 6 | 6 | **5** |

In Figure 5, we show rootograms for the MD-NB (top row) and MD-ZINB (bottom row). We see that the MD-NB under-fits count 0 and over-fits the subsequent counts 1 and 2 in the first component, which is typical for data with excess zeros. The rootograms for the MD-ZINB fit the data better in the first component. Rootograms nicely exhibit the different means and variances for each component.

Component 1 contains only 6% (430/7118) of all observations and corresponds to individuals who spend, on average, fewer days in hospital but with very high variance. The mode of hospital days for component 1 is 4 for the MD-NB and 7 for the MD-ZINB (zero mode omitted). Component 2 is the largest, with 59% (4172/7118) of individuals; they stay longer in hospital (the mode is at 24 days) with less variance. Component 3 comprises 35% (2517/7118) of the population, and these patients have the most hospital days (mode at 39 days) and again high variance.

Figure 3 shows *β*_*k*_ parameter estimates for each component of the MD-NB as incidence rate ratios (IRRs) alongside the Bayesian high probability density intervals. These coefficients can be interpreted as the multiplicative increase in the expected number of hospital days for every one unit increase in the predictor. For example, in components 1 and 2, treatment is associated with more hospital days, across all modalities. The IRR for the combination of all three treatments is the largest, 7.9. That is, compared to patients who receive no treatment, those who receive chemotherapy, radiation, and surgery have 7.9 times as many expected hospital days. In general, all treatment combinations have higher IRRs in component 1 than in components 2 and 3. The only exception is radiotherapy, which has an IRR of 4.1 in component 2, higher than 0.6 in component 3, and 3.8 in component 1. The IRR for the number of metastases is uneven over the components: 1.24 and 1.22 in 1 (MD-NB and MD-ZINB), 1.51 in 2, and 0.95 in 3. On the other hand, the IRR for the number of multiple tumors is increasing from 0.69 to 0.91 to 0.94 across components 1, 2, and 3. In component 3, radiation is associated with fewer hospital days and surgery is null, whereas chemotherapy and combinations are associated with more hospital days. Demographic factors and baseline health were less strongly associated with hospital days. Age and sex appear to have no relationship to hospital days (the IRRs are around 1.0 in all components). Regional factors are only important for individuals in component 3 and only for urban districts, where the IRR is 0.64.

**Figure 3:**
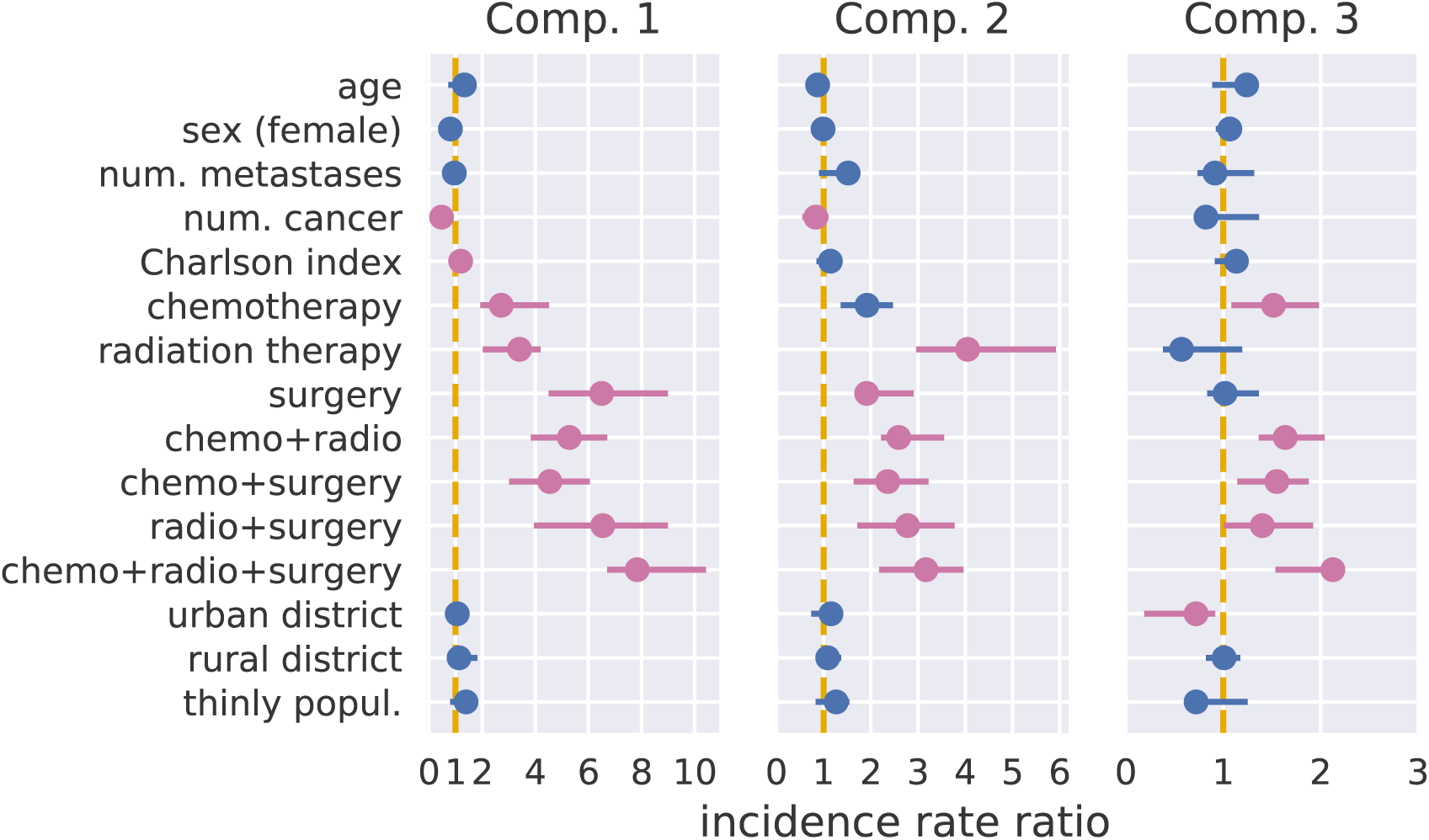
NP-NB estimation results for all three components on the AOK data set. Parameter estimates are presented as incidence rate ratios and 95% highest probability density intervals. Intervals that exclude the 1 are highlighted in purple. Intercept is not shown.

We display the regression coefficient estimates for component 1 of the MD-ZINB in Figure 4, separately for the negative binomial part of the model (left) and the binomial part of the model (right). Note that the binomial coefficients are relative risks, while the negative binomial coefficients are IRRs. In the negative binomial component, the pattern of coefficients is nearly the same as the MD-NB fit, but the coefficients are slightly different. For example, the IRR for receiving all three treatments (chemotherapy, radiotherapy, and surgery) is 7.6 in the NB components of the MD-ZINB compared to 7.9 in the MD-NB. We do not show regression coefficient estimates for components 2 and 3 of the MD-ZINB because they are essentially the same as the MD-NB coefficients.

**Figure 4:**
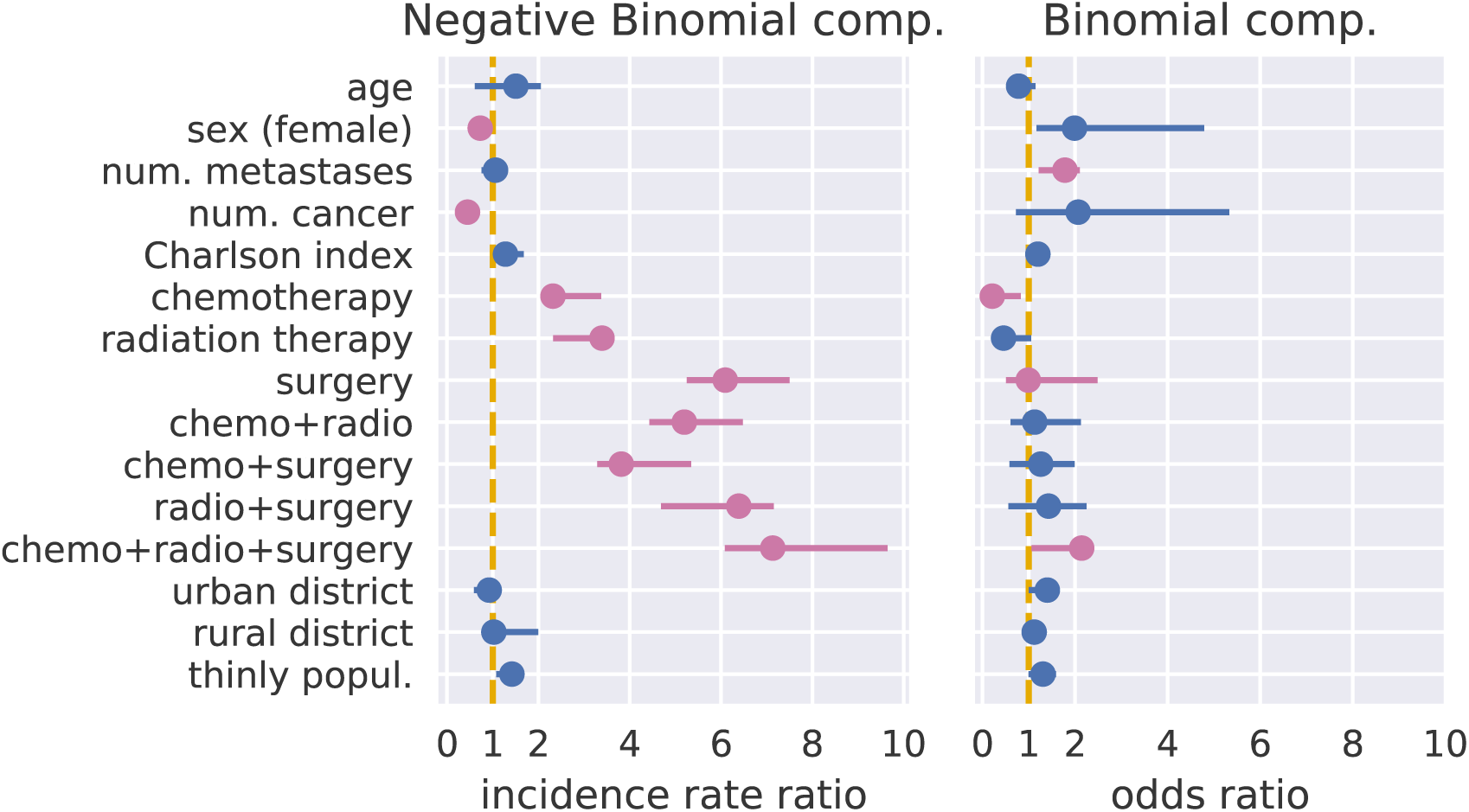
NP-ZINB estimation results for the first components only on the AOK data set. Parameter estimates are presented as incidence rate ratios for the count part and odds ratios for the zero part. 95% highest probability density intervals are shown for both parts. Intervals that exclude the 1 are highlighted in purple. Intercept is not shown.

When we use hard assignments to classify individuals into components according to the highest posterior probability, we see that treatment patterns are very different across components (see Figure 6). Chemotherapy plus radiation is the most common treatment in all components, but individuals in component 1 are far more likely to receive this combination (42% compared to 22% in component 2 and 24% in component 3). Surgery alone is the second most common treatment in components 2 (22%) and 3 (17%), and chemotherapy alone is the second most common treatment in component 1 (19%), but it is infrequent in 2 (13%) and 3 (11%). Compared to people in component 1, those in components 2 and 3 are more likely to receive chemotherapy combined with surgery or surgery and radiation. Figure 9 shows greater comorbidity burden and slightly older age in component 3. The Charlson comorbidity burden increased across the components: median 1 (IRQ 0–3) in component 1; median 2 (IQR 0–3) in component 2; and median 2 (IQR 1–4) in component 3. Metastases were more common in patients in component 1 (median 1, IQR 0–1) than in components 2 and 3 (median 0, IQR 0–1).

**Figure 5:**
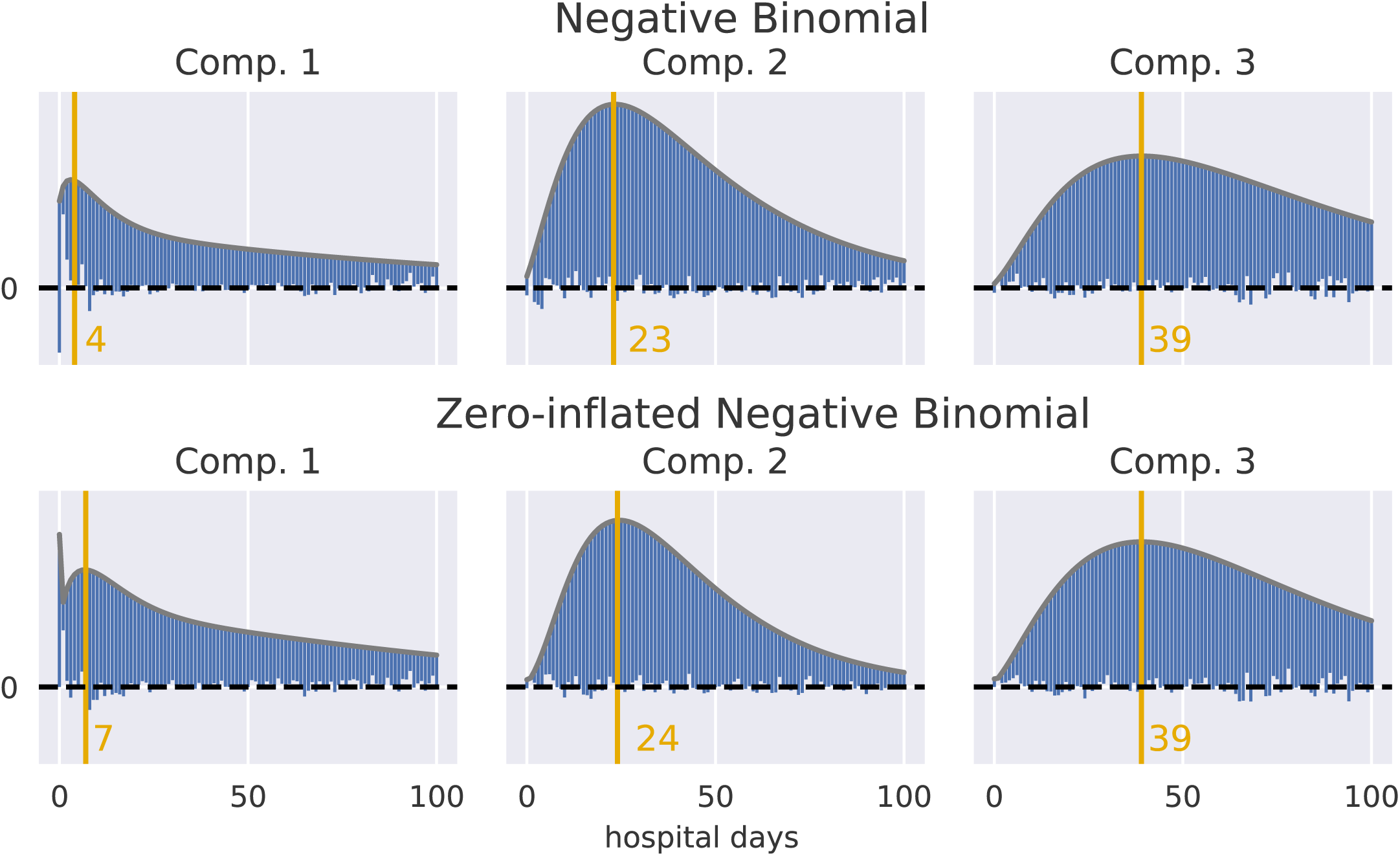
Hanging rootograms showing goodness-of-fit by comparing observed frequencies from the model to expected frequencies for NP-NB (top row) and NP-ZINB (bottom row) for all three components based on component assignment weights from the posterior. Ideal fit is indicated when the bars do not overlap or underlie the horizontal zero line. The vertical line marks the mode.

**Figure 6:**
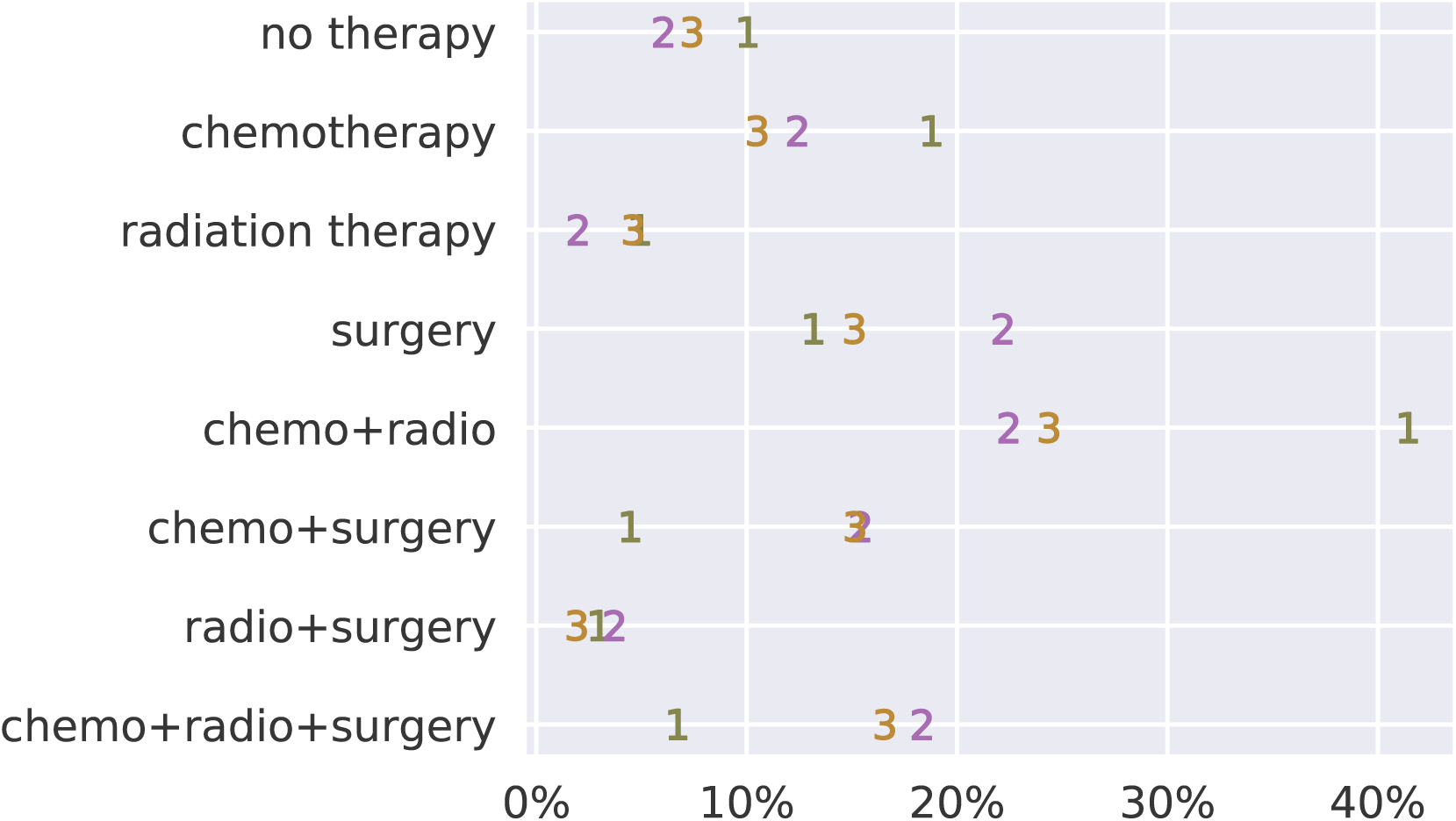
Plot for treatment types as percentages after using hard component assignments based on the MD-ZINB for the AOK data set. Numbers mark the three components.

**Figure 7:**
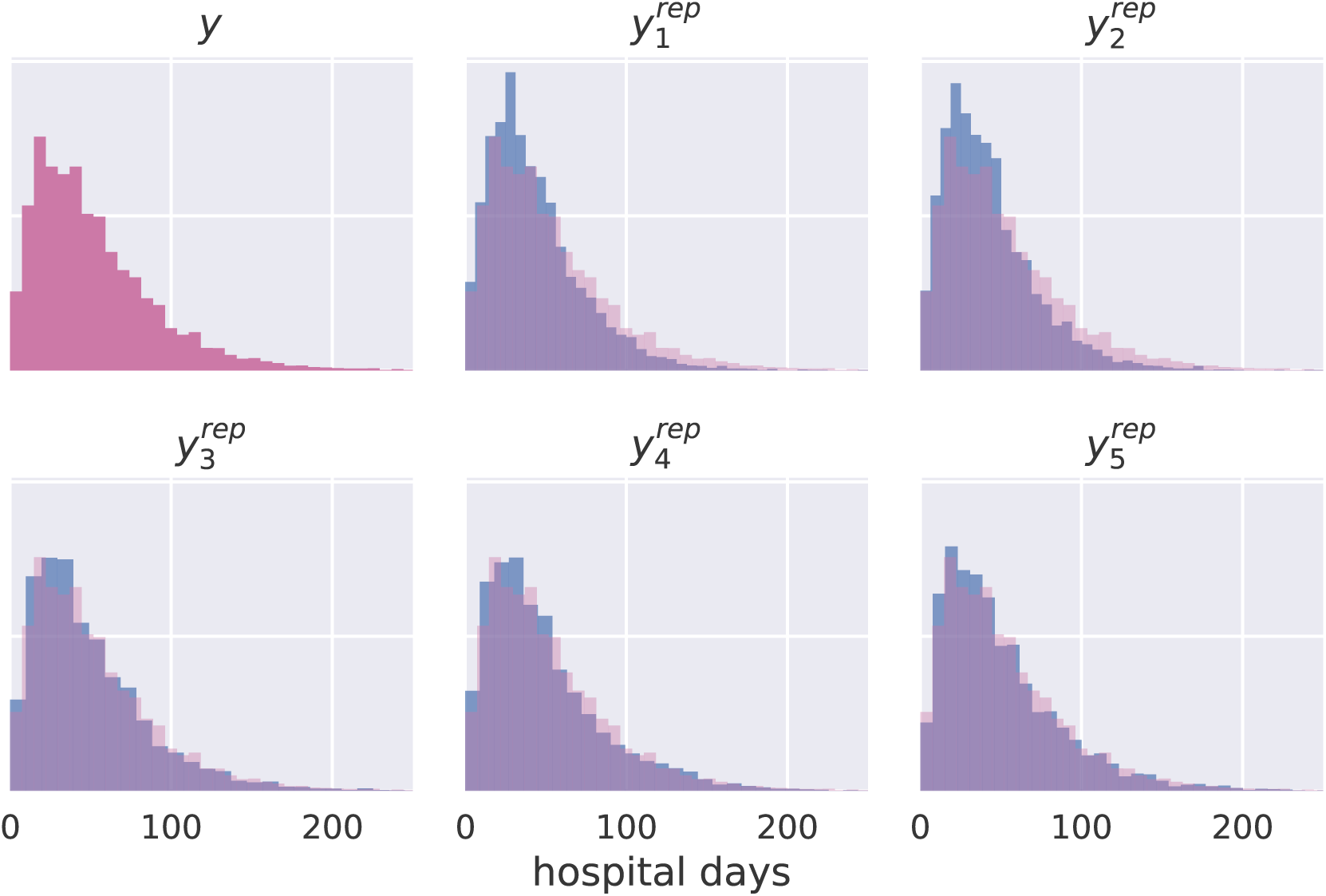
Density histograms for replicated outcome *y*^*rep*^ from simulating data from the posterior predictive distribution using the observed predictors. True outcome *y* from the AOK data set for comparison.

**Figure 8:**
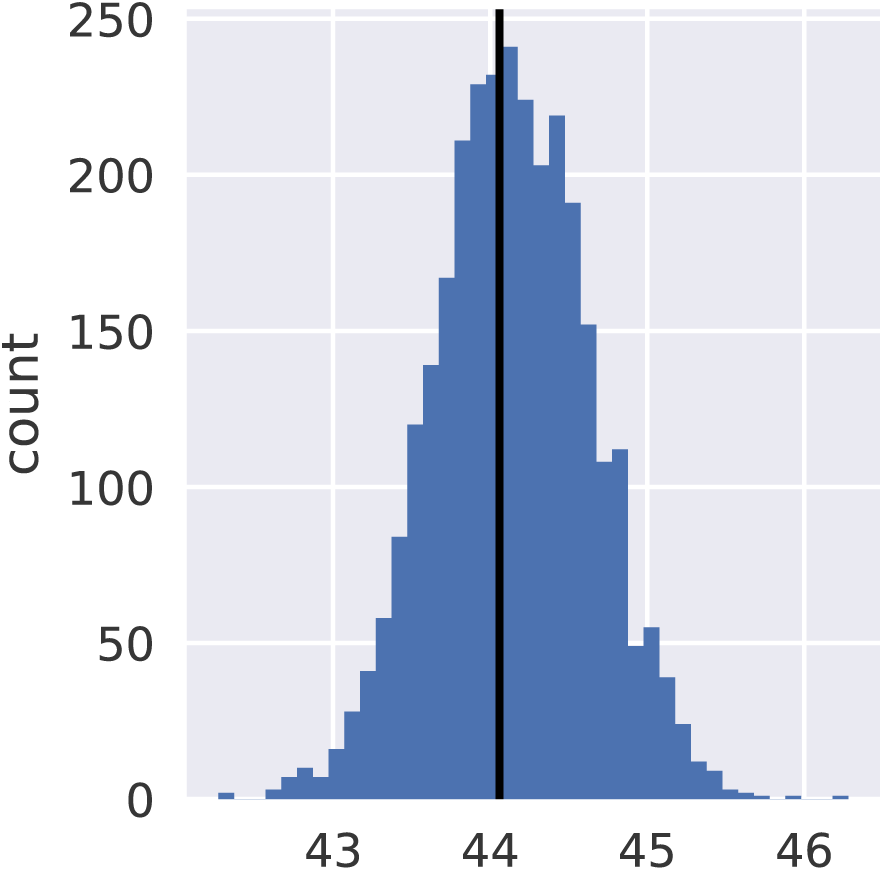
Histogram of means based on 1000 replicated data sets from the posterior predictive distribution. The black line marks the observed mean of hospital days from the AOK data set.

**Figure 9:**
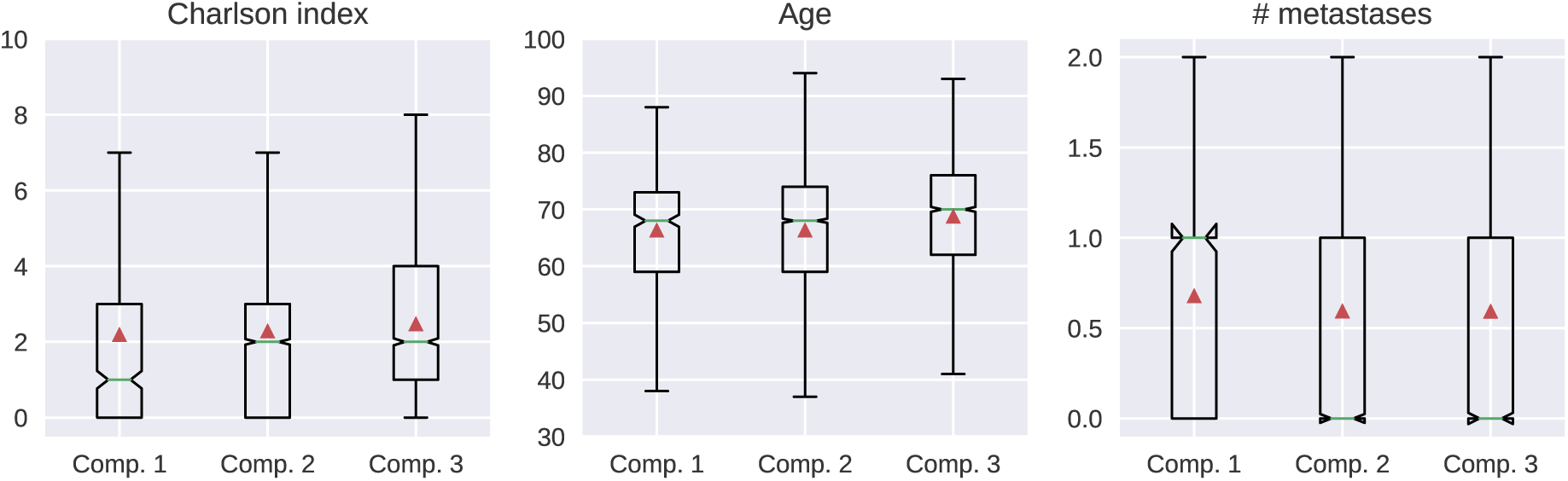
Boxplots for Charlson comorbidity index, age, and the number of metastases after using hard component assignments based on the MD-ZINB for the AOK data set. The red triangle marks the mean.

## 6 Discussion

This paper compares parametric Bayesian models and finite mixture models for count data on health care utilization. The advantage of Bayesian mixture modeling is to allow the number of mixture components to be estimated from the data. In a simulation study, we show that selecting the number of mixture components in a finite mixture model using model fit statistics such as AIC and BIC is not a very accurate method for finding the true number of mixture components. Instead, the posterior mixture component probabilities from the Bayesian model are closer to the truth, though slightly overestimate the number of components, as seen by other authors. [60]

However, our simulation only covered a small fraction of possible scenarios. Furthermore, the data-generating process closely resembles the model specification, which is rarely the case in the real world. While the MD-NB is clearly more accurate in finding the true number of components than AIC or BIC model selection, it still misses the exact components in many of the simulations. In this simulation, “truth” is defined by the data-generating process, but it has been argued that the idea of “truth” in component analysis depends on the context and the application. [61]

On the AOK data set, graphical model checks show that the model fits the data well, particularly the zero-inflated model. This is not clear from the histograms, as there is no apparent spike at zero, but becomes obvious in the rootograms. For the AOK data set, the MD-NB and MD-ZINB models find three components of individuals with strikingly different distributions of hospital days and treatment patterns.

In the treatment of lung cancer, surgery offers the best prospect of cure. If diagnosed at an early stage, it is possible to remove the tumor as a whole, such that no further treatment is necessary. If the tumor is already bigger, surgical resection with chemotherapy and radiation is the treatment of choice. In metastatic lung cancer, palliative chemotherapy, possibly accompanied by radiation therapy for individual metastases, may alleviate symptoms and prolong survival. [62]

Component 1 has the fewest hospital days on average. In this component, we find many patients with chemotherapy only, and chemotherapy in combination radiation therapy. This, and the lack of surgery, likely indicate that these patients were already in an advanced (metastatic) stage at diagnosis. For these patients, it is likely that therapy had a palliative intent with a focus on improving quality of life. In contrast, patients in components 2 and 3 were more likely to have surgery only, surgery and chemotherapy, and the combination of all three treatments. This indicates diagnosis at an earlier stage and more aggressive treatment.

The treatments received by people in components 2 and 3 are quite similar, with only the proportion of surgery being slightly higher in 2 than in 3. However, the coefficients governing the relationships between treatment type and hospital days in these components are quite different. While radiation therapy is associated with significantly more hospital days in component 2, it has the opposite association in component 3. Moreover, the strong and positive association of surgery with hospital days in component 2 fades in component 3, where surgery has no relationship to hospital days. Together, these results suggest that people in components 2 and 3 get similar treatment combinations, but for different reasons.

There are several limitations to this study. First, mixture models present computational challenges. For example, care must be taken when fitting Bayesian mixture models to avoid the so-called “label-switching problem” caused by the model being invariant under permutations of the indices of the components (i.e., the indices of the model components may be permuted across chains). [33, 63] It is crucial to run multiple Markov chains, inspect the resulting posterior samples, and apply posterior checks, as we have done here. We also enforced an ordering constraint on the component means. Other authors have proposed more sophisticated methods, using loss functions, [64, 65] exploratory analysis of unconstrained posterior samples from a permutation sampler, [13] or highest posterior density. [66] The number of selected components might vary, as mixture models with different values of *K* can provide good representations of the same data. [67] Our choice of *K* = 20 balances computational burden with the the goal of our analysis, which is to describe the parameters of the mixture components and the corresponding clusters of observations. More than 20 components/clusters would be unwieldy in our applied setting. [68] The appropriate limit on the number of mixture components would be different in a mixture model intended for flexible density estimation, for which the approximation to the infinite mixture occurs at values of *K* nearer to 70 or 100. [69–71] In our Bayesian model, components that contribute very little will have mixing proportions that go to zero, thus being effectively removed from the model. This allows us to make a single training run in which we start with a relatively large initial value of *K*, and allow surplus components to be pruned out of the model. [32]

In our applications, fitting this model to the AOK data took from approximately 4 hours up to 12 hours, depending on the number of observations, on a current quadcore CPU with 32GB RAM. The finite mixture model with a pre-specified number of components took only 3 minutes to compute. Further research should investigate how variational Bayesian methods could improve speed and how this affects the accuracy of the estimates. Variational Bayesian mixture models have been found to accurately detect the number of mixture components [67, 72] and suffer less from identifiability issues. [73]

We limited our consideration to mixtures of parametric (zero-inflated) Negative Binomial regression models. Previous authors have recommended more flexible kernels for count data. [74, 75] The ZINB distribution is more flexible than the Poisson, but still parametric. [37] suggested a semi-parametric alternative to two-part models for zero-inflated data.

We are also limited by the parametric form of the mixing distribution. [76] proposed a semiparametric approach to modeling flexibly the relationship of covariates to mixing proportions. There exists an extensive literature on *nonparametric* Bayesian mixture modeling approaches, (see reviews in [77–80] and references therein). The Dirichlet prior is the most widely used prior for mixture components, [81, 82] though [74] found the Pitman–Yor process can be more robust. [83] notes that the Dirichlet distribution is limited the negative correlation structures and proposes to use the Beta-Liouville distribution. Relevant to our application, [84] find that Bayesian nonparametric mixture models of medical claims outperform analogous mixture models, though their purpose was prediction rather than interpretation and they modeled semicontinuous, rather than zero-inflated count, outcomes. [85] noted the connection between mixtures of Poisson and negative binomial and Dirichlet processes.

Some authors have argued that interpreting the parameters of the mixture coefficients should be avoided. [71, 86] In addition, interpretation is necessarily more difficult in complex models such as these. In the case of the MD-ZINB model, with three mixture components in each of two sub-models (i.e., the Binomial part and the Negative Binomial part), the number of regression coefficients is six times the number of covariates (assuming each covariate is in each sub-model). However, inference on multiple parameters simultaneously is relatively straightforward in Bayesian models, which is another advantage of this approach.

## 7 Conclusion

This work presents Bayesian and likelihood-based clustering mixture models for count data with many zeros (here, hospital days) that can be used to find subgroups of patients. In contrast to clustering methods based on finite mixture models, the Bayesian approach avoids under- and over-fitting while still being fully interpretable. We apply this method to study hospital days for patients with lung cancer and demonstrate that it can find subgroups with specific properties that correspond well to the different number of hospital days in each component. Clustering models are useful and practical methods for understanding heterogeneity in inpatient hospital services, an important component of total health care spending.

## 8 Acknowledgments

The authors thank Bret Zeldow for reviewing the manuscript, Michael Betancourt and Bob Carpenter for help with the Stan code, and Larissa Schwarzkopf for interpreting the clusters.

## Appendix

**Table 2:** Data-generating parameter for the simulation study. N indicates the number of draws in each component.

| Component | N | high overlap |  |  |  | medium overlap |  |  |  | low overlap |  |  |  |
| --- | --- | --- | --- | --- | --- | --- | --- | --- | --- | --- | --- | --- | --- |
| | | $\beta_0$ | $\beta_1$ | $\beta_2$ | $\psi$ | $\beta_0$ | $\beta_1$ | $\beta_2$ | $\psi$ | $\beta_0$ | $\beta_1$ | $\beta_2$ | $\psi$ |
| 1 | 3000 | 0.0 | 0.5 | -0.5 | 1.00 | 0.0 | 0.5 | -0.5 | 1.00 | 0.0 | 0.5 | -0.5 | 1.00 |
| 2 | 5000 | 1.0 | 0.0 | 0.0 | 13.25 | 1.8 | 0.0 | 0.0 | 25.75 | 3.2 | 0.0 | 0.0 | 50.75 |
| 3 | 3000 | 1.7 | -0.5 | 0.5 | 25.50 | 2.4 | -0.5 | 0.5 | 50.50 | 3.9 | -0.5 | 0.5 | 100.50 |
| 4 | 4000 | 2.0 | 0.0 | 0.0 | 37.75 | 2.7 | 0.0 | 0.0 | 75.25 | 4.3 | 0.0 | 0.0 | 150.25 |
| 5 | 2000 | 2.3 | 0.5 | -0.5 | 50.00 | 3.0 | 0.5 | -0.5 | 100.00 | 4.6 | 0.5 | -0.5 | 200.00 |

**Table 3:**
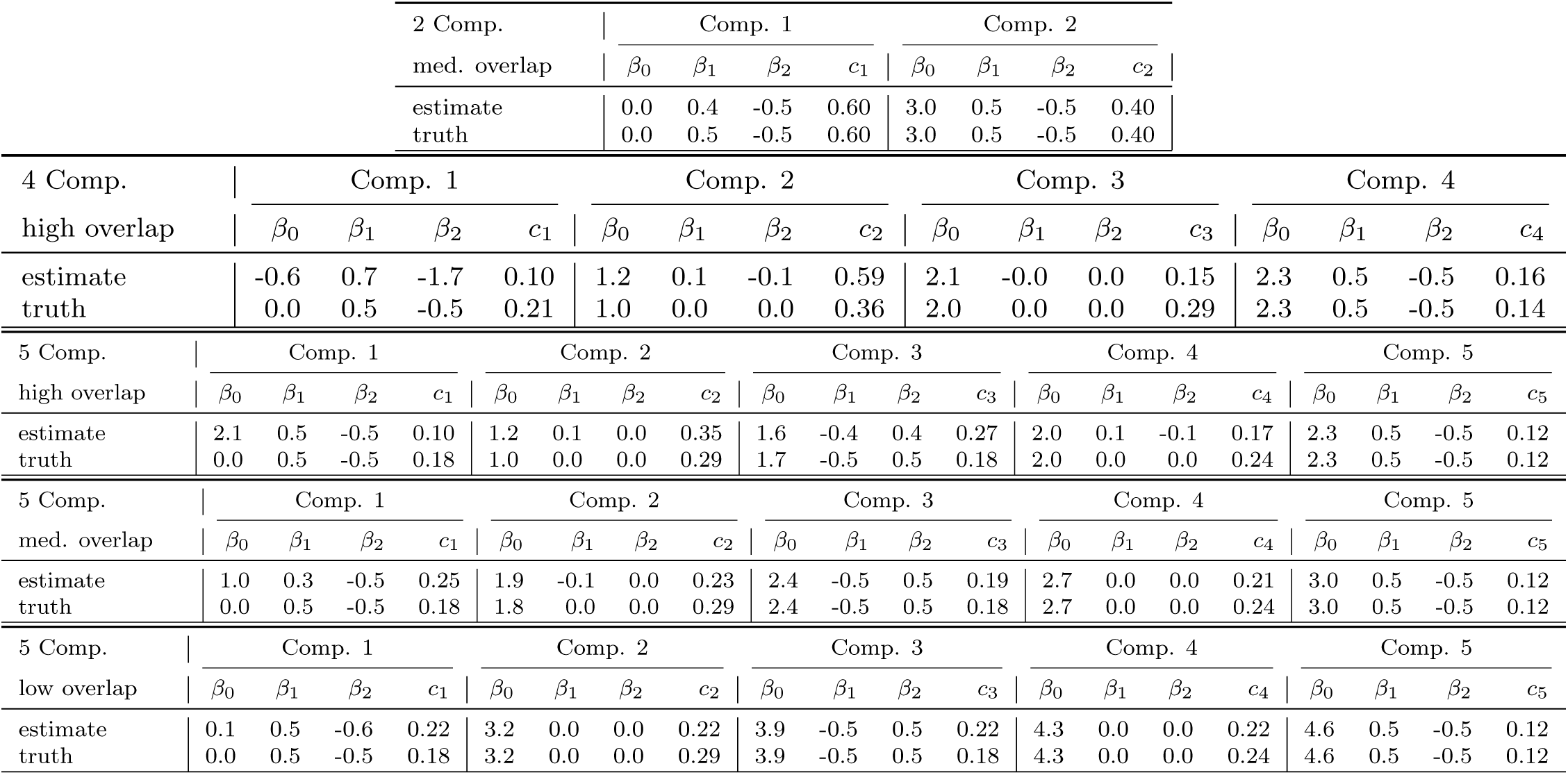
Comparison of true and estimated parameters for cases where the number of mixture components was accurately estimated by MD-NB. *β*’s indicate intercept and regression coefficients, *c*’s are the mixture weights. All estimates are based on posterior medians. Because the estimated mixture components are not always in the same order, estimated components were hand-matched to their corresponding best-fitting component.

| 2 Comp. |  | Comp. 1 |  |  |  | Comp. 2 |  |  |  |
| --- | --- | --- | --- | --- | --- | --- | --- | --- | --- |
| med. overlap | | $\beta_0$ | $\beta_1$ | $\beta_2$ | $c_1$ | $\beta_0$ | $\beta_1$ | $\beta_2$ | $c_2$ |
| estimate |  | 0.0 | 0.4 | -0.5 | 0.60 | 3.0 | 0.5 | -0.5 | 0.40 |
| truth |  | 0.0 | 0.5 | -0.5 | 0.60 | 3.0 | 0.5 | -0.5 | 0.40 |

| 4 Comp. |  | Comp. 1 |  |  |  | Comp. 2 |  |  |  | Comp. 3 |  |  |  | Comp. 4 |  |  |  |
| --- | --- | --- | --- | --- | --- | --- | --- | --- | --- | --- | --- | --- | --- | --- | --- | --- | --- |
| high overlap | | $\beta_0$ | $\beta_1$ | $\beta_2$ | $c_1$ | $\beta_0$ | $\beta_1$ | $\beta_2$ | $c_2$ | $\beta_0$ | $\beta_1$ | $\beta_2$ | $c_3$ | $\beta_0$ | $\beta_1$ | $\beta_2$ | $c_4$ |
| estimate |  | -0.6 | 0.7 | -1.7 | 0.10 | 1.2 | 0.1 | -0.1 | 0.59 | 2.1 | -0.0 | 0.0 | 0.15 | 2.3 | 0.5 | -0.5 | 0.16 |
| truth |  | 0.0 | 0.5 | -0.5 | 0.21 | 1.0 | 0.0 | 0.0 | 0.36 | 2.0 | 0.0 | 0.0 | 0.29 | 2.3 | 0.5 | -0.5 | 0.14 |

Table 3: Comparison of true and estimated parameters for cases where the number of mixture components was accurately estimated by MD-NB.
| 5 Comp. |  | Comp. 1 |  |  |  | Comp. 2 |  |  |  | Comp. 3 |  |  |  | Comp. 4 |  |  |  | Comp. 5 |  |  |  |
| --- | --- | --- | --- | --- | --- | --- | --- | --- | --- | --- | --- | --- | --- | --- | --- | --- | --- | --- | --- | --- | --- |
| high overlap | | $\beta_0$ | $\beta_1$ | $\beta_2$ | $c_1$ | $\beta_0$ | $\beta_1$ | $\beta_2$ | $c_2$ | $\beta_0$ | $\beta_1$ | $\beta_2$ | $c_3$ | $\beta_0$ | $\beta_1$ | $\beta_2$ | $c_4$ | $\beta_0$ | $\beta_1$ | $\beta_2$ | $c_5$ |
| estimate |  | 2.1 | 0.5 | -0.5 | 0.10 | 1.2 | 0.1 | 0.0 | 0.35 | 1.6 | -0.4 | 0.4 | 0.27 | 2.0 | 0.1 | -0.1 | 0.17 | 2.3 | 0.5 | -0.5 | 0.12 |
| truth |  | 0.0 | 0.5 | -0.5 | 0.18 | 1.0 | 0.0 | 0.0 | 0.29 | 1.7 | -0.5 | 0.5 | 0.18 | 2.0 | 0.0 | 0.0 | 0.24 | 2.3 | 0.5 | -0.5 | 0.12 |
| med. overlap | | $\beta_0$ | $\beta_1$ | $\beta_2$ | $c_1$ | $\beta_0$ | $\beta_1$ | $\beta_2$ | $c_2$ | $\beta_0$ | $\beta_1$ | $\beta_2$ | $c_3$ | $\beta_0$ | $\beta_1$ | $\beta_2$ | $c_4$ | $\beta_0$ | $\beta_1$ | $\beta_2$ | $c_5$ |
| estimate |  | 1.0 | 0.3 | -0.5 | 0.25 | 1.9 | -0.1 | 0.0 | 0.23 | 2.4 | -0.5 | 0.5 | 0.19 | 2.7 | 0.0 | 0.0 | 0.21 | 3.0 | 0.5 | -0.5 | 0.12 |
| truth |  | 0.0 | 0.5 | -0.5 | 0.18 | 1.8 | 0.0 | 0.0 | 0.29 | 2.4 | -0.5 | 0.5 | 0.18 | 2.7 | 0.0 | 0.0 | 0.24 | 3.0 | 0.5 | -0.5 | 0.12 |
| low overlap | | $\beta_0$ | $\beta_1$ | $\beta_2$ | $c_1$ | $\beta_0$ | $\beta_1$ | $\beta_2$ | $c_2$ | $\beta_0$ | $\beta_1$ | $\beta_2$ | $c_3$ | $\beta_0$ | $\beta_1$ | $\beta_2$ | $c_4$ | $\beta_0$ | $\beta_1$ | $\beta_2$ | $c_5$ |
| estimate |  | 0.1 | 0.5 | -0.6 | 0.22 | 3.2 | 0.0 | 0.0 | 0.22 | 3.9 | -0.5 | 0.5 | 0.22 | 4.3 | 0.0 | 0.0 | 0.22 | 4.6 | 0.5 | -0.5 | 0.12 |
| truth |  | 0.0 | 0.5 | -0.5 | 0.18 | 3.2 | 0.0 | 0.0 | 0.29 | 3.9 | -0.5 | 0.5 | 0.18 | 4.3 | 0.0 | 0.0 | 0.24 | 4.6 | 0.5 | -0.5 | 0.12 |

**Table 4:**
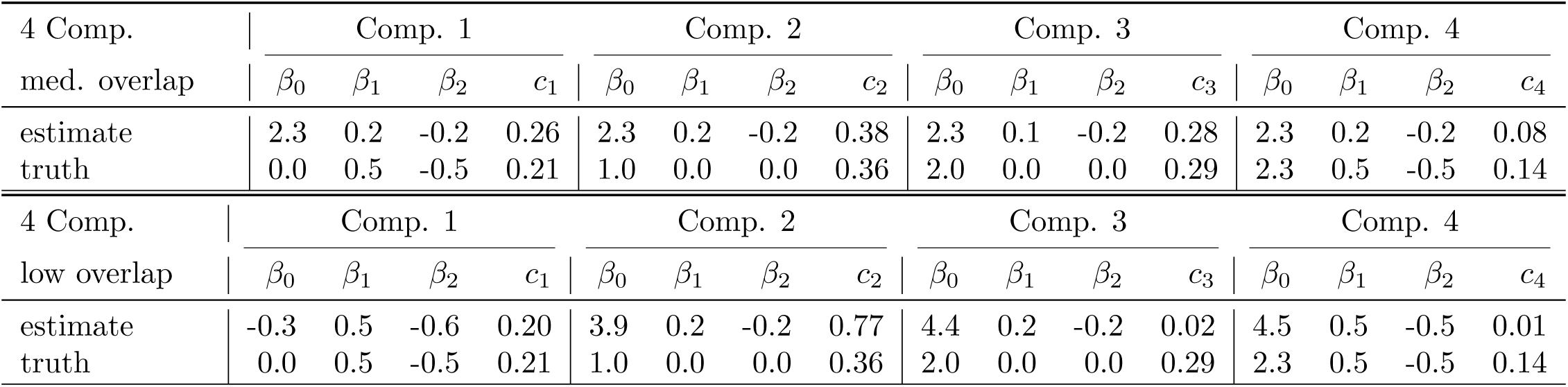
Comparison of true and estimated parameters for cases where the number of mixture components was accurately estimated by AIC and BIC. *β*’s indicate intercept and regression coefficients, *c*’s are the mixture weights.

## Footnotes

1 Throughout, we use “component” to refer to the individual distributions in the mixture and “cluster” to denote the observations drawn from each mixture component.

